# Combined single-cell gene and isoform expression analysis in haematopoietic stem and progenitor cells

**DOI:** 10.1101/2020.04.06.027474

**Authors:** Laura Mincarelli, Vladimir Uzun, Stuart A. Rushworth, Wilfried Haerty, Iain C. Macaulay

## Abstract

Single-cell RNA sequencing (scRNA-seq) enables gene expression profiling and characterization of novel cell types within heterogeneous cell populations. However, most approaches cannot detect alternatively spliced transcripts, which can profoundly shape cell phenotype by generating functionally distinct proteins from the same gene. Here, we integrate short- and long-read scRNA-seq of hematopoietic stem and progenitor cells to characterize changes in cell type abundance, gene and isoform expression during differentiation and ageing.

## Main text

Single-cell RNA-seq (scRNA-seq) technologies are now applied to a broad spectrum of biological systems ^1^ with particular impact in the study of stem cell and developmental biology ^2–4^. With the advance of short-read technologies capable of analysing many thousands of cells in a single experiment, it has become possible to identify cell types and changes in complex biological systems, such as the haematopoietic hierarchy.

Alternative splicing (AS) is thought to occur in at least 62% of multi-exonic genes in mouse (Gencode M24^5^) and up to 95% of multi-exonic genes in human^6^ representing a key means for the generation of additional proteome diversity. Given that most short-read scRNA-seq approaches target just a short region of a transcript, typically at the 3’ or 5’ ends, AS is unlikely to be captured and an entire class of information about cell- and cell-type specific AS is overlooked.

Here, we have integrated parallel short- (Illumina) and long-read (Pacific Biosciences; PacBio) sequencing of single-cell libraries to profile cellular diversity, gene expression and AS events in the mouse haematopoietic system. We isolated the **L**ineage negative, c**K**it/Cd117 positive (**LK**) cell fraction - containing haematopoietic stem and progenitor cells ^7^ from young (8 week) and aged (72 week) C57/BL6 mice by Fluorescence Activated Cell Sorting (FACS). We generated cell-barcoded cDNA using the 10X Genomics Chromium platform and prepared both PacBio libraries from this full-length cDNA, as well as standard Illumina libraries (**Fig 1A, Supplementary Fig 1**)

**Figure 1:**
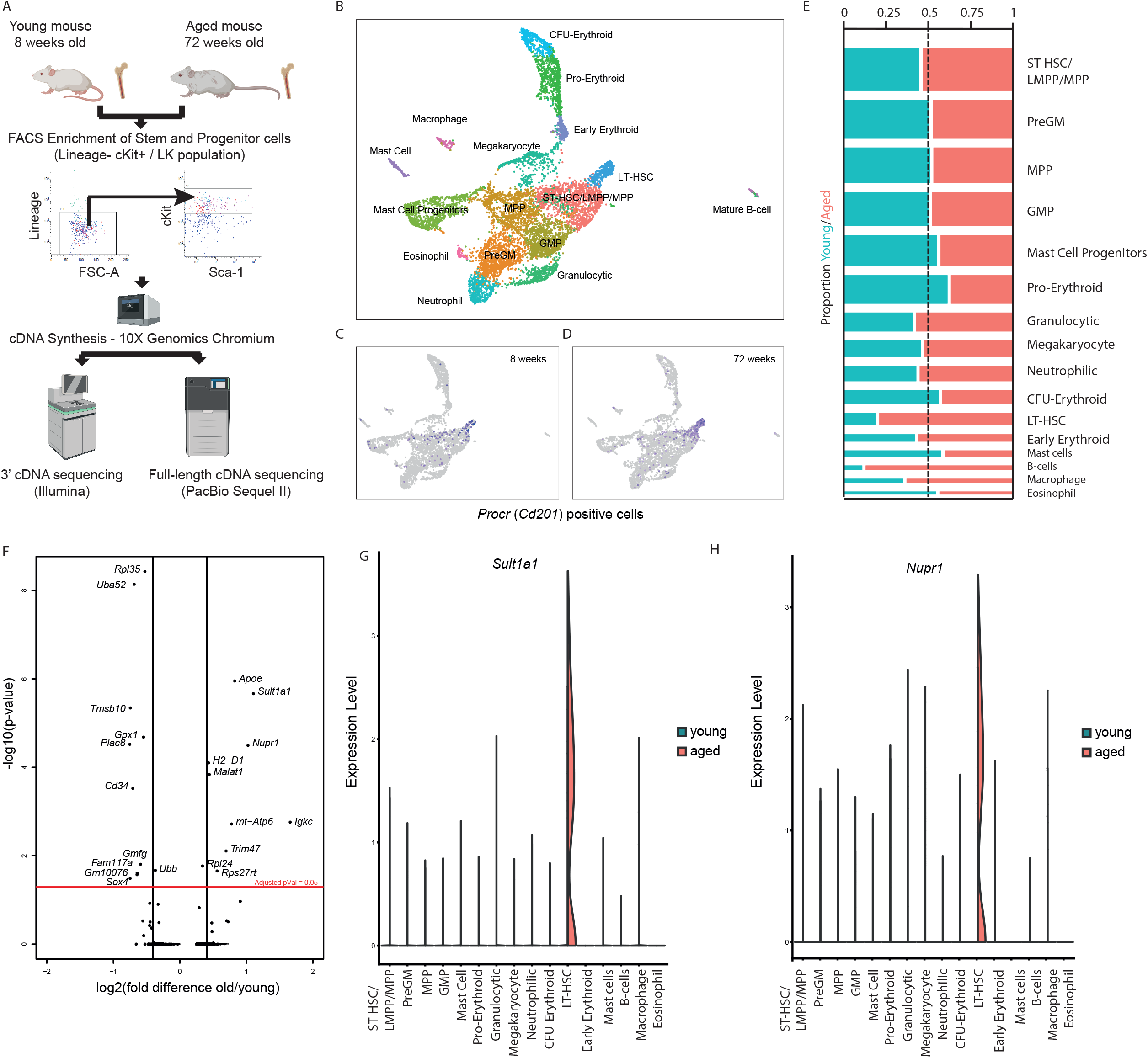
Profiling changes in cellular composition and gene expression in ageing haematopoiesis. **A**) Experimental overview. Following sorting, Lin− cKit+ cells were analysed by parallel short- and long- read scRNA-seq. **B**) Cell clustering revealed cellular heterogeneity in the Lin− cKit+ cell population, including a distinct population of Procr+ LT-HSCs in both young and old mice (**C**, **D**), which was expanded in the aged mice (**E**). **F**) Differential gene expression between young and aged LT-HSCs identified a number of differentially expressed genes, including the LT-HSC specific genes *Sult1a1* (**G**) and *Nupr1* (**H**).

The 10X/Illumina sequencing data revealed the diversity of cell types present within the 8,000 LK cells passing quality control (**Fig 1B**) correlating with the diversity expected from phenotypic analysis of the same population ^7^. Overall, our analysis identified 15 subclusters within the LK population, including long-term haematopoietic stem cells (LT-HSCs), here associated with *Procr* (*Cd201*) expression^8^, as well as intermediate and committed progenitor cells (**Fig 1B**). Small numbers of mature B-cells, myeloid cells and mast cells were also observed, but were transcriptionally very distinct from the main stem and progenitor cell cluster, and most likely represent a low level of circulating cells. While the majority of cell type abundances were unchanged between 8- and 72-week old mice, there was an increased abundance of phenotypic LT-HSCs (**Fig 1C-E**) in keeping with previous findings based on stem cells defined by protein marker expression (Lineage negative, Sca-1 positive, cKit positive (LSK) CD34− CD48− CD150+ cells)^9^.

To identify a transcriptional signature of LT-HSC ageing, we compared short-read gene expression levels specifically in the population of young and aged HSCs. We observed upregulation of several genes including *Sult1a1*, *Nupr1*, *ApoE* and *Igkc* and downregulation of *Rpl35*, *Uba52*, *Tmsb10*, *Gpx1*, *Plac8* and *Cd34* (**Fig 1F, Supplementary Table 1**). *Sult1a1* and *Nupr1* are highly restricted to the LT-HSC population, and are indeed almost exclusively expressed in aged LT-HSCs (**Fig 1G-H**).

The genes encoding *Nupr1* and *Sult1a1* are co-localised in a 50 kb region of chromosome 7, and their elevated expression has previously been shown to be associated with an age-related increase in the H3K4me3 chromatin mark in HSCs^10^. *Sult1a1* encodes for a sulfotransferase which acts on substrates including hormones and neurotransmitters, and *Nupr1* has a regulatory role in cell proliferation and apoptosis, but neither has a described functional role in haematopoiesis ^10,11^. The age-associated upregulation of the immunoglobulin kappa constant region *Igkc* in LT-HSCs was unexpected, but not without precedent. Previous work has shown that *Igkc* transcripts are expressed in aged LT-HSCs, possibly as a result of epigenetic dysregulation ^12^. Thus, this short-read scRNA-seq analysis of young and aged stem and progenitor cells can identify changes in cell abundance and gene expression associated with ageing, and highlights potential novel markers of the aged LT-HSC population.

In order to further examine transcriptional heterogeneity in the haematopoietic system, we performed long-read PacBio sequencing (IsoSeq) on the cDNA pools generated from the 10X platform, similar to an approach recently applied in cerebellar cells^13^. This protocol, taking advantage of the cell barcoding technology used in 10X Genomics library preparation, enables isoform identification and association with cell populations, and potentially even single-cells, through integration of the long- and short- read data.

A detailed breakdown of the PacBio sequencing statistics is presented in **Supplementary Table 2**. In brief, PacBio sequencing yielded a total of 17.9 million circular consensus sequencing (CCS) reads with a median read length of 1471 bases (**Fig 2A**). These reads mapped to an average of 16,427 genes per sample, representing an average of 33,345 transcripts per sample and an average of 31 reads per transcript. Transcript coverage averaged 74% (**Fig 2B**), and alternative isoforms were detected for 52.3% of genes(**Fig 2C**). The combined dataset further revealed 2820 previously undescribed exons present in 2165 genes supported by at least two PacBio reads (**Supplementary Table 3**).

**Figure 2:**
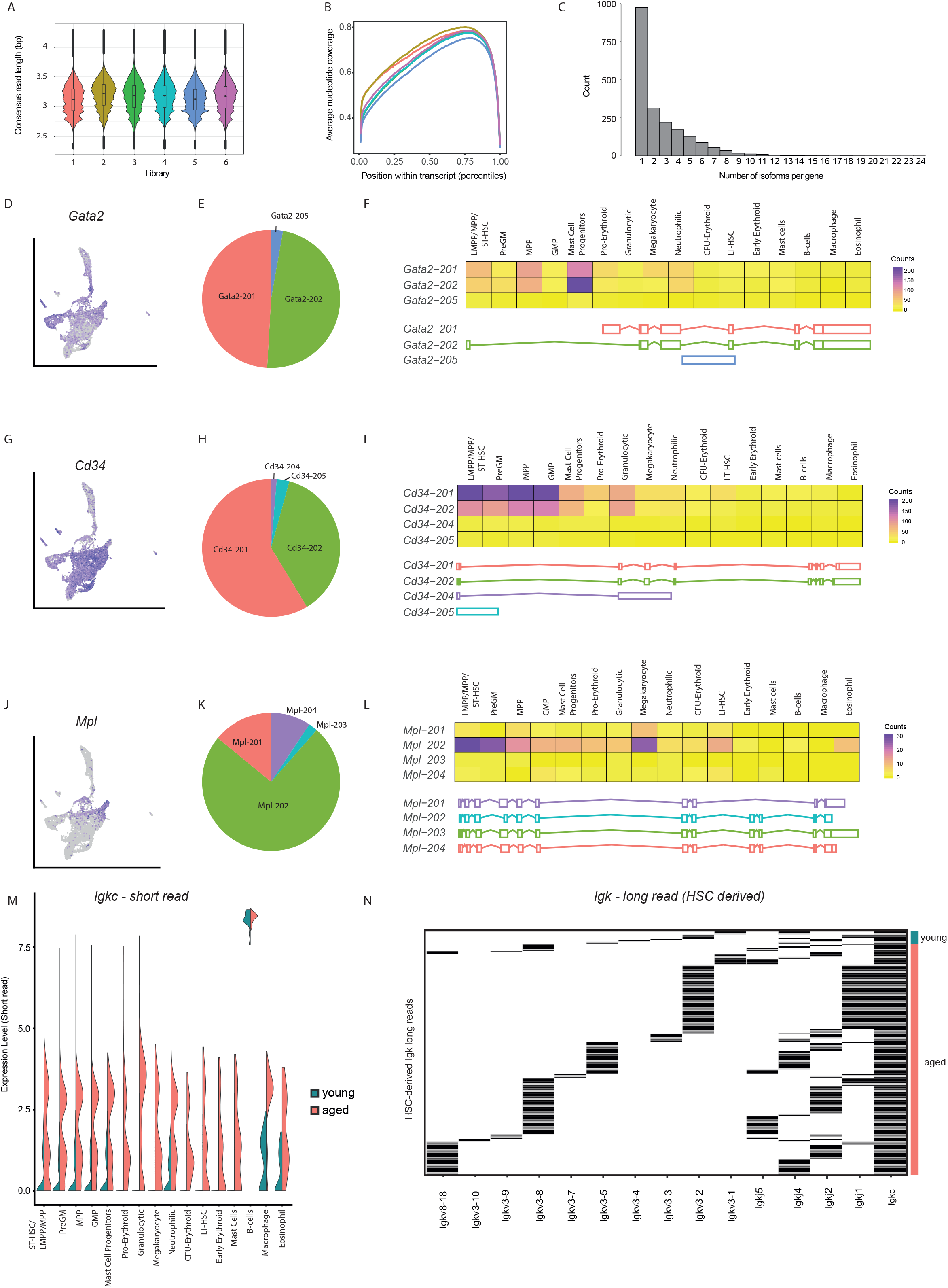
Integration of long- and short- read data to profile cell type isoform usage. **A**) Read length distribution from long-read sequencing of 10X single-cell libraries. **B**) transcript coverage from long-read sequencing of 10X single-cell libraries. **C**) Number of isoforms detected per gene. **D**) short-read expression of *Gata2*, **E**) distribution of *Gata2* isoforms in the demultiplexed data and **F**) cell type distribution of reads for the detected *Gata2* isforms. **G**,**H**,**I**) and **J**,**K**,**L**) as above for *Cd34* and *Mpl*. **M**) Upregulation of *Igkc* throughout the hematopoietic system in aged mice **N**) Detection of recombined *Igkc* reads in LT-HSCs.

Demultiplexing of the long reads using the cell barcodes identified in the short-read analysis enabled 5.8 million (32%) of CCS reads to be assigned to individual cells. Using the multi-modal capabilities of Seurat^14^, we integrated short- and long- read datasets allowing side-by-side comparison of gene and isoform expression from the respective platforms, and annotation of individual cells with isoform-level transcript information. A median of 411 reads, corresponding to a median of 292 isoforms, could be assigned per cell (**Supplementary Fig 2A and 2B**). In general, the numbers of isoforms detected are too low to allow meaningful comparisons at single-cell resolution. However, with 50-500,000 reads per cluster, scaling linearly with the number of cells per cluster (**Supplementary Fig 2C-E**), cell-type expression can be assessed.

AS was observed for a number of key haematopoietic genes, including *Gata2, Cd34* and *Mpl*, each of which had multiple detectable isoforms (**Fig 2D-K**). These examples highlight the impact of additional long-read sequencing in scRNA-seq studies; where short-read analysis alone would have predicted the expression of single isoforms, by integrating long reads we demonstrate that multiple variants are commonly expressed. These variants resulting from splicing events are either internal to the gene models (e.g. *Mpl*) or at the 5’ end of the genes (e.g. *Gata2*), and would thus have been undetectable by short reads alone.

*Gata2* expression, which in our short read data is restricted to stem and progenitor cell, mast cell and early megakaryocyte and erythroid progenitor populations (**Fig 2D**), consists of two main isoforms, *Gata2-201* and *Gata2-202* (**Fig 2E**) – which are translated into the same protein but are differentiated by the usage of distinct distal exons. The *Gata2-202* isoform is specific to haematopoietic and neuronal lineages, while the *Gata2-201* isoform shows less specificity in its expression ^15^. Here, both isoforms are most abundant in the mast cell population, with *Gata2-202* showing more specific expression in this cell type (**Fig 2F**).

*Cd34* expression is restricted to progenitor cells, and is largely absent in the HSC, MK/E and mature mast/myeloid populations (**Fig 2G**). The long-read sequencing data reveals that two isoforms predominate, *Cd34-201* and *Cd34-202* (**Fig 2H and I**), the latter encoding a truncated variant lacking the majority of its cytoplasmic tail which has been speculated to have a functional impact on cell signalling ^16,17^.

Three predominant isoforms of *Mpl*, the transcript encoding the thrombopoietin receptor ^18^, were detectable in both stem cells and megakaryocytes (**Fig 2J-L**). The primary isoform, *Mpl-201* encodes the transmembrane Mpl receptor. *Mpl-202* encodes a shortened version of the protein, lacking 8 amino acids in the extracellular domain. *Mpl-204*, which lacks exons 9 and 10 completely encodes a truncated protein with no transmembrane domain. This isoform is detectable in the stem cell population, and functional studies have indicated that it has an inhibitory role on normal Mpl signalling^18^.

Following the observation that IgK transcripts were upregulated in the aged LT-HSC population, and indeed throughout the myeloid progenitor (**Fig 2M**), we used the long read data to determine that the IgK arising from the LT-HSC population in aged mice consisted not just of “germline” IgK, as previously reported^12^, but fully VDJ-recombined transcripts **(Fig 2N, Supplementary Fig 3)**. While unprecedented, there is an increasing body of evidence that the expression of recombined Ig molecules in non-lymphoid cells is possible ^19–21^, often in age-related diseases such as acute myeloid leukemia^22^.

Thus, long-read sequencing can readily add complementary data and uncover additional heterogeneity within scRNA-seq studies. AS and isoform expression are currently largely overlooked in scRNA-seq studies in spite of their immense functional significance in shaping cellular phenotype. Here, we demonstrate that long-read sequencing can be integrated into conventional single-cell approaches to predict functional diversity at the proteomic level. We demonstrate that over half of the genes detected in a standard scRNA-seq analysis have multiple isoforms expressed and that functionally distinct isoforms of critical regulatory genes are prevalent.

With continued improvements in the throughput of long-read sequencing platforms and development of methods targeting specific cells or transcripts, we envision that long-read sequencing will enable detailed characterisation of isoform diversity at the single-cell level at a scale comparable to current short-read based methods. This will be applicable, for example, to large-scale “atlassing” studies of cellular heterogeneity, and in haematological malignancies where mutations in splicing factors are common ^23^.

## Contributions

LM, SAR and ICM designed and performed experiments. LM, VU, WH and ICM analysed data.

## Methods

### Stem and Progenitor cell isolation

All animal work in this study was carried out in accordance with regulations set by the United Kingdom Home Office and the Animal Scientific Procedures Act of 1986. Bone marrow was isolated from spine, femora, tibiae, and ilia of 8 weeks and 72 weeks old C57BL/6 mice. Red blood cell depletion was performed with ammonium chloride lysis (STEMCELL Technologies) and lineage negative cells were isolated using the EasySep Mouse Hematopoietic Progenitor Cell Isolation Kit (STEMCELL Technologies).

The lineage depleted cells were stained with following fluorophore-conjugated monoclonal antibodies: Cd105-PE, clone MJ7/18, Miltenyi; Cd4-Vioblue, clone REA604, Miltenyi; Cd11b-Vioblue, clone REA592, Miltenyi; Cd117-Pe Vio770, clone REA791, Miltenyi; Cd8a-Vioblue, clone 53-6.7, Miltenyi; CD50-Vioblue, clone REA421, Miltenyi; CD45R-Vioblue, clone REA755, Miltenyi; GR1-Vioblue, clone REA810, Miltenyi; Sca-APC, clone REA422, Miltenyi; CD48-APC Cy7, clone HM48-1, Miltenyi; CD150-BV510, clone TC15-12F12, CD34-PeCy5, MEC147, Miltenyi. Approximately 10,000 LK (Lin-, CD117+) cells per sample were sorted using the BD FACSMelody cell sorter (BD Biosciences, San Jose, California) into 1 x PBS containing 5% BSA.

### Sequencing of single-cell cDNA libraries

Sorted cells were processed by 3’ end single cell RNA-Seq using the 10X Genomics Chromium (V2 Kit) according to manufacturer’s protocol (10X Genomics, Pleasanton, CA) with an increase to 16 cycles for the cDNA PCR amplification. Libraries were sequenced on a NextSeq500 (Illumina, San Diego) in paired end, single index mode as per the 10X recommended metrics.

Raw Illumina sequencing data were analysed with the 10X Genomics CellRanger pipeline (version 3.0.2) to obtain a single-cell expression matrix object. Subsequent analysis was performed in R using Seurat version 3 ^14^. Cells showing gene counts lower than 1,000 and a mitochondrial gene expression percentage higher than 5% were excluded from further analysis. Within Seurat, data were normalised using NormalizeData (normalization.method = “LogNormalize”, scale.factor = 10000) and data from multiple samples were merged using the FindIntegrationAnchors and IntegrateData commands.

### Pacific Biosciences Sequel Sequencing of single-cell cDNA libraries

Libraries compatible the Pacific Biosciences Sequel/Sequel II systems were prepared from 800 ng input cDNA, following the “No size selection” Iso-Seq library preparation according to the manufacturer’s instructions (IsoSeq Template Preparation for Sequel System V05), with the following modifications: The elution incubation time during AMPure beads purification was increased to 10 minutes and the second AMPure bead purification step, following the exonuclease reaction, was omitted to optimise library concentration.

### Pacific Biosystems long read analysis

Circular Consensus reads (CCS) were generated using the following parameters: maximum subread length 20,000, minimum subread length 50 and minimum number of passes 3.

Reads with identified polydA or polydT were demultiplexed using bbduk https://sourceforge.net/projects/bbmap/) (k = 16, hdist = 3) using the 10X genomics barcodes identified from the short-read analysis. Long reads were mapped to the mouse genome (mm10) using Minimap 2 (v2.17), and to the gencode (vM19) transcriptome.

Novel exons were identified by investigating the alignment of the reads to the transcriptome identifying inserts of at least 21 nucleotides located at exon junctions. To confirm the existence of these exons, the alignment of the reads to the genome was parsed, exonic sequences located within the previously identified intron, and supported by at least to reads were retained for further analysis. We further removed any exonic regions overlapping RefSeq annotations (GRCm38, last accessed February 19 2019).

To identify reads supporting V(D)J recombination events, we used IgBlast v1.14.0 ^24^) using default parameters (-min_V_length 9 -min_J_length 0 -min_D_match 5 -D_penalty -2 - J_penalty -2).

### Data integration in Seurat

In order to reduce the batch effect, count matrices produced by short read sequencing for individual libraries were combined in Seurat using FindIntegrationAnchors and IntegrateData functions (dims = 1:20).

Illumina and PacBio reads were integrated in Seurat using the CreateAssayObject command to add the long read data to an existing Seurat object already containing the short read data. This links the demultiplexed long-reads with the short-read data through the cell barcodes present in both.

## Supplementary Figures

**Supplementary Figure 1: Experimental overview.** Full-length cDNA is generated by the 10X Genomics Chromium instrument. This material is used as input for conventional short-read (Illumina) and long-read (PacBio) sequencing, which enables sequencing of the full-length of the transcript and detection of alternative splicing or recombination events. Cell clustering is performed on the short-read data. Integrating the cell barcodes present in both the short- and long-read data enables assignment of long-reads to cells and cell clusters.

**Supplementary Figure 2: Long-read sequencing from 10X Genomics Chromium cDNA A**) Number of long reads per cell, separated by cluster (as identified in Fig 1B) **B**) Number of transcripts per cell, again separated by cluster **C**) Total number of long-reads per cluster **D**) Total number of transcripts per cluster. **E**) Correlation of number of long-reads per cluster with cluster cell number.

**Supplementary Figure 3: Reads from the Immunoglobulin kappa locus.** Reads from young and aged LT-HSCs visualised in the Integrated Genome Viewer (IGV) show recombination between the Igkc locus and the J domains.

## Supplementary Tables

**Supplementary Table 1: Differential gene expression between young and aged HSCs (related to Figure 1F)**.

**Supplementary Table 2: Long-read sequencing statistics**

**Supplementary Table 3: Novel Exons detected in Long-read sequencing data.**

