## Supplementary figures and images for "Combined single-cell gene and isoform expression analysis in haematopoietic stem and progenitor cells"

### Supplementary Figure 1

Supplementary Figure 1

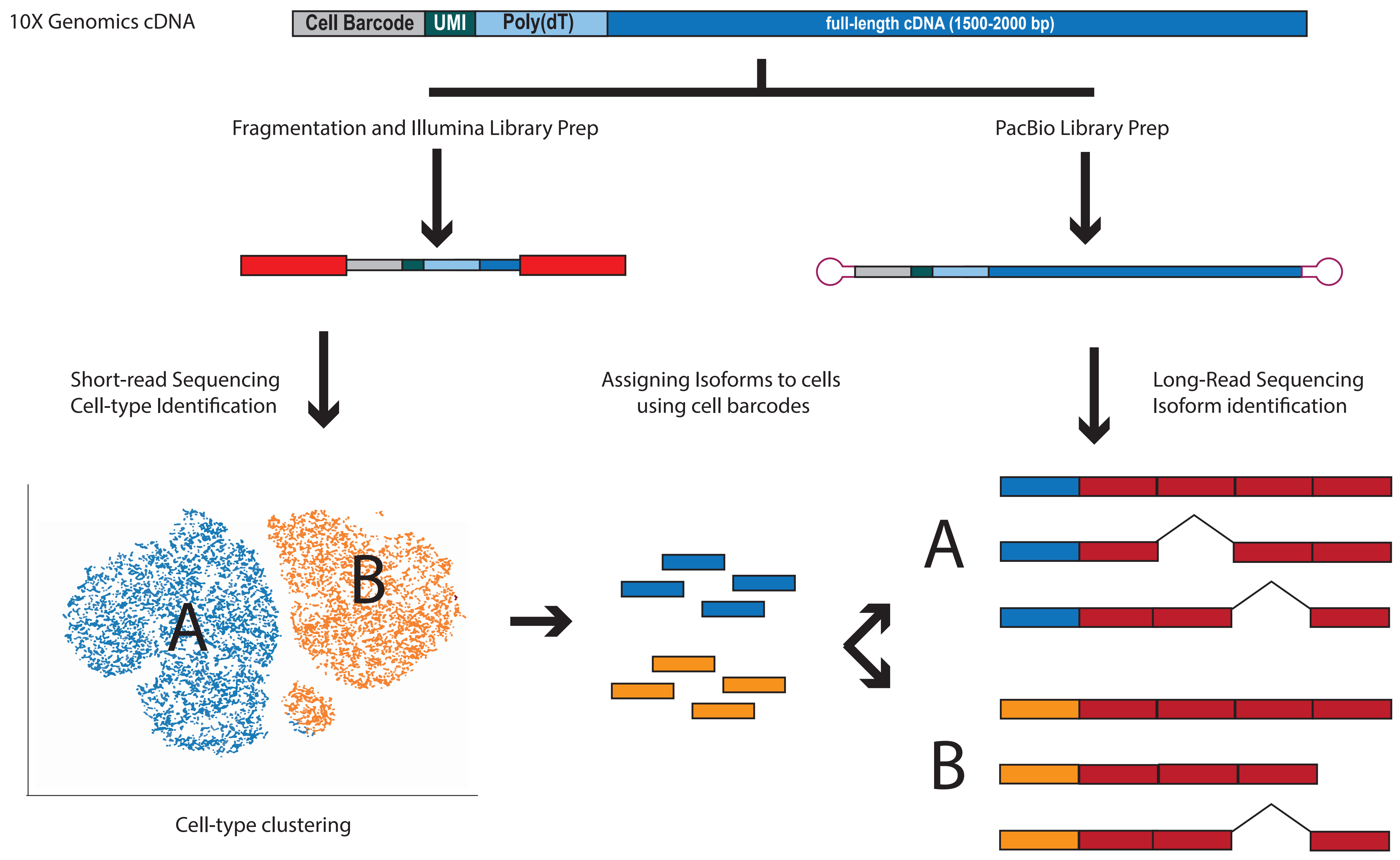

### Supplementary Figure 2

Supplementary Figure 2

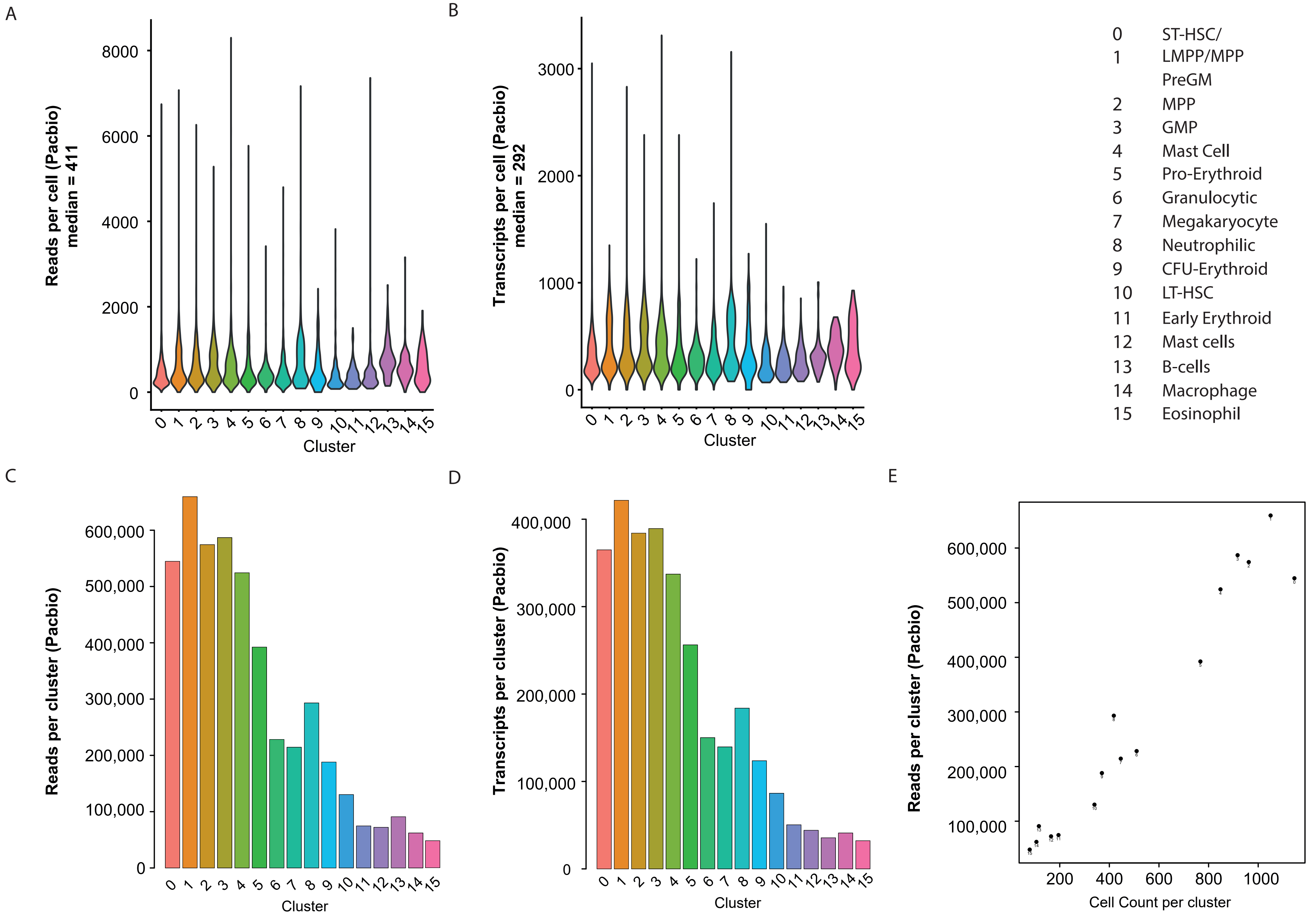

### Supplementary Figure 3

#### Supplementary Figure 3

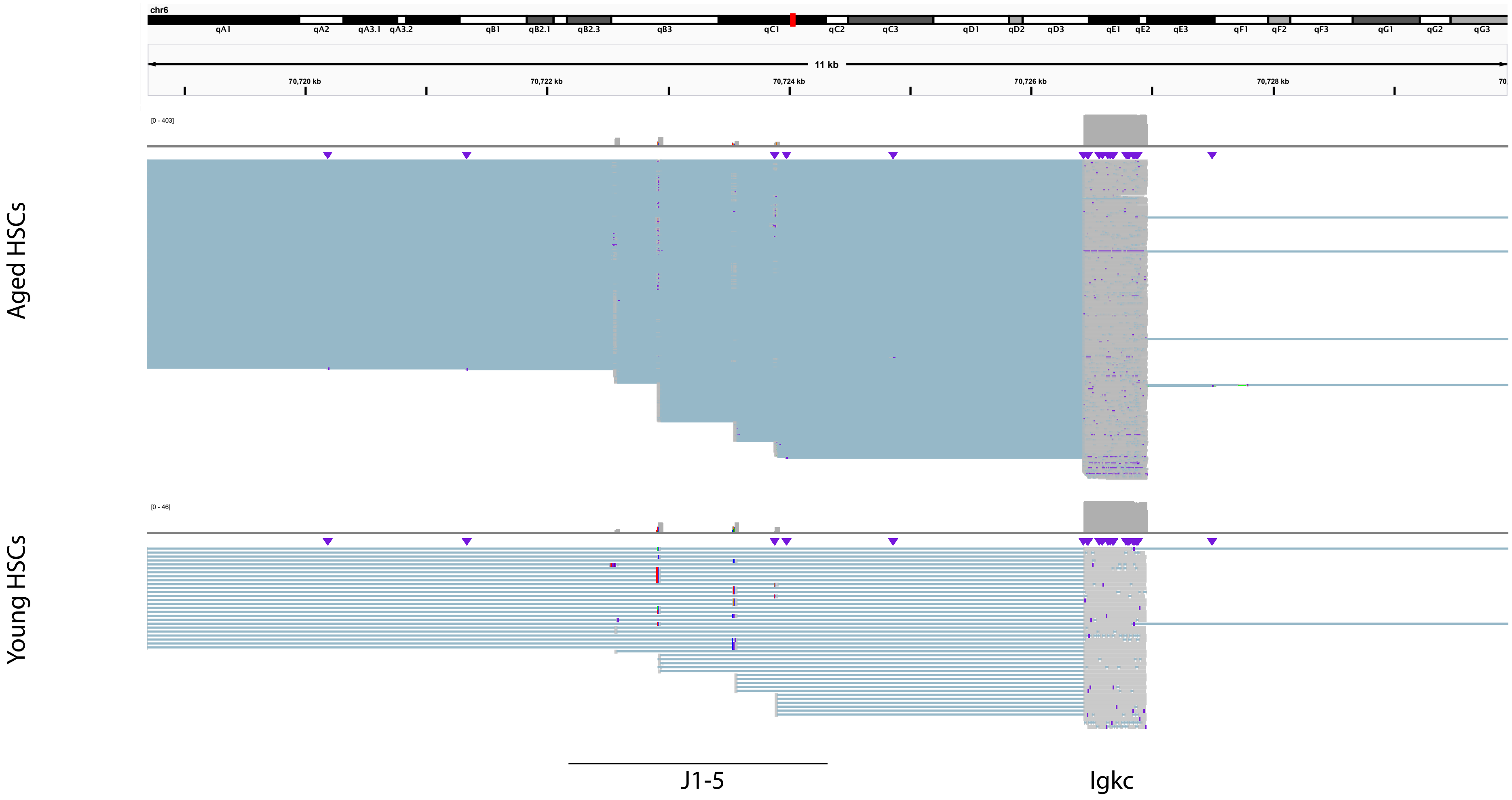
